# The expected time to cross extended fitness plateaus

**DOI:** 10.1101/343053

**Authors:** Mahan Ghafari, Daniel Weissman

## Abstract

For a population to acquire a complex adaptation requiring multiple individually neutral mutations, it must cross a plateau in the fitness landscape. We consider plateaus involving three mutations, and show that large populations can cross them rapidly via lineages that acquire multiple mutations while remaining at low frequency. Plateau-crossing is fastest for very large populations. At intermediate population sizes, recombination can greatly accelerate adaptation by combining independent mutant lineages to form triple-mutants. For more frequent recombination, such that the population is kept near linkage equilibrium, we extend our analysis to find simple expressions for the expected time to cross plateaus of arbitrary width.

## 1 Introduction

Most mutations in most natural populations are effectively neutral. Considered in isolation, these are irrelevant for adaptation. But the fitness effect of a mutation generally depends on the genetic background on which it occurs, a phenomenon known as epistasis. Thus, there are likely to be combinations of these neutral mutations that interact epistatically to have an effect on fitness. If this effect is positive for a given combination, then that combination forms a *complex adaptation*, separated from the wild type by a fitness plateau. How frequently do we expect populations to acquire such adaptations? On one hand, a given complex adaptation should typically be harder for a population to find than a simple adaptation requiring only a single beneficial mutation. On the other hand, if a genome of length *L* has 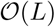 possible neutral mutations, then there are 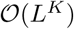 genotypes that could potentially be a complex adaptation involving *K* mutations. So if even a modest fraction of these genotypes are indeed adaptive, the number of possible complex adaptations could far exceed the number of available beneficial mutations, and it could be that they are collectively a frequent form of adaptation [Fisher, 2007, Weissman et al., 2009, Trotter et al., 2014]. To evaluate their importance, we must know more about how rapidly population explore fitness plateaus.

Populations can cross fitness plateaus via a sequence of neutral mutations fixing by drift until only one more mutation is needed for the (formerly complex) adaptation. But this process is slow and inefficient; in a high-dimensional fitness plateau, the population will be much more likely to drift away from a complex adaptation than towards it. Large, asexual populations can cross plateaus and even fitness valleys much more rapidly (e.g., [van Nimwegen and Crutchfield, 2000, Komarova et al., 2003, Iwasa et al., 2004, Weinreich and Chao, 2005, Durrett and Schmidt, 2008, Weissman et al., 2009]). They can do this because many mutations will be present in the population at low frequency. If the population is sufficiently large, even these low-frequency mutations will be present in a large absolute number of individuals, some of which will happen to also carry additional mutations. Thus, genotypes that are multiple mutations away from the consensus genotype will already be present in the population and exposed to natural selection, allowing the population to effectively “see” several steps away in the fitness landscape, and “tunnel” directly to the adaptive genotypes Jain and Krug [2006].

Recombination changes these dynamics in two ways. First, by combining mutations that occur in different lineages, it accelerates the population’s exploration of the plateau Christiansen et al. [1998]. On the other hand, recombination breaks up the beneficial combination once it is formed Eshel and Feldman [1970], Feldman [1971], Karlin and McGregor [1971], slowing adaptation Takahata [1982], Michalakis and Slatkin [1996]. While the latter effect is fairly easy to understand quantitatively, the former depends on the spectrum of mutant lineages that coexist in the population and has only been fully understood in the simplest case of two-locus plateaus [Weissman et al., 2010]. Here we extend this analysis to the three-locus case, considering the full spectrum of possible population sizes, recombination rates, mutation rates, and selective advantages of the adaptive genotype. We also analyze the dynamics for arbitrary-width plateaus when recombination is frequent relative to selection.

## 2 Model

We consider a haploid Wright-Fisher population of size *N.* The genome consists of *K* loci each of which has two possible alleles, 0 and 1; for much of the analysis, we will focus on the case *K* = 3. Initially, all individuals have the all-0 genotype. All genotypes have the same fitness except the all-1 genotype, which has a strong selective advantage *s* ≫ 1/*N*; see Figure 1. Individuals mutate (in both directions) at a rate µ per locus per generation. Each generation, each offspring is produced clonally (with possible mutations) with probability 1 − *r*; with probability *r*, it is the product of recombination between two parents. Recombinant offspring sample each locus independently with equal probability from their parents (again, with possible mutation). We will focus on finding the expected time 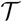 until the all-1 genotype first makes up the majority of the population. For simplicity, we will also refer to the “rate” of crossing the plateau, defined as 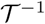, even though it is not a true rate, as the distribution of time to cross the plateau is not exponential in general. The definitions of the most important symbols are collected in Table 1. Exact simulations were done in Python (Figs. 3 and 4) and Mathematica (Figs. 5 and 6).

**Figure 1:**
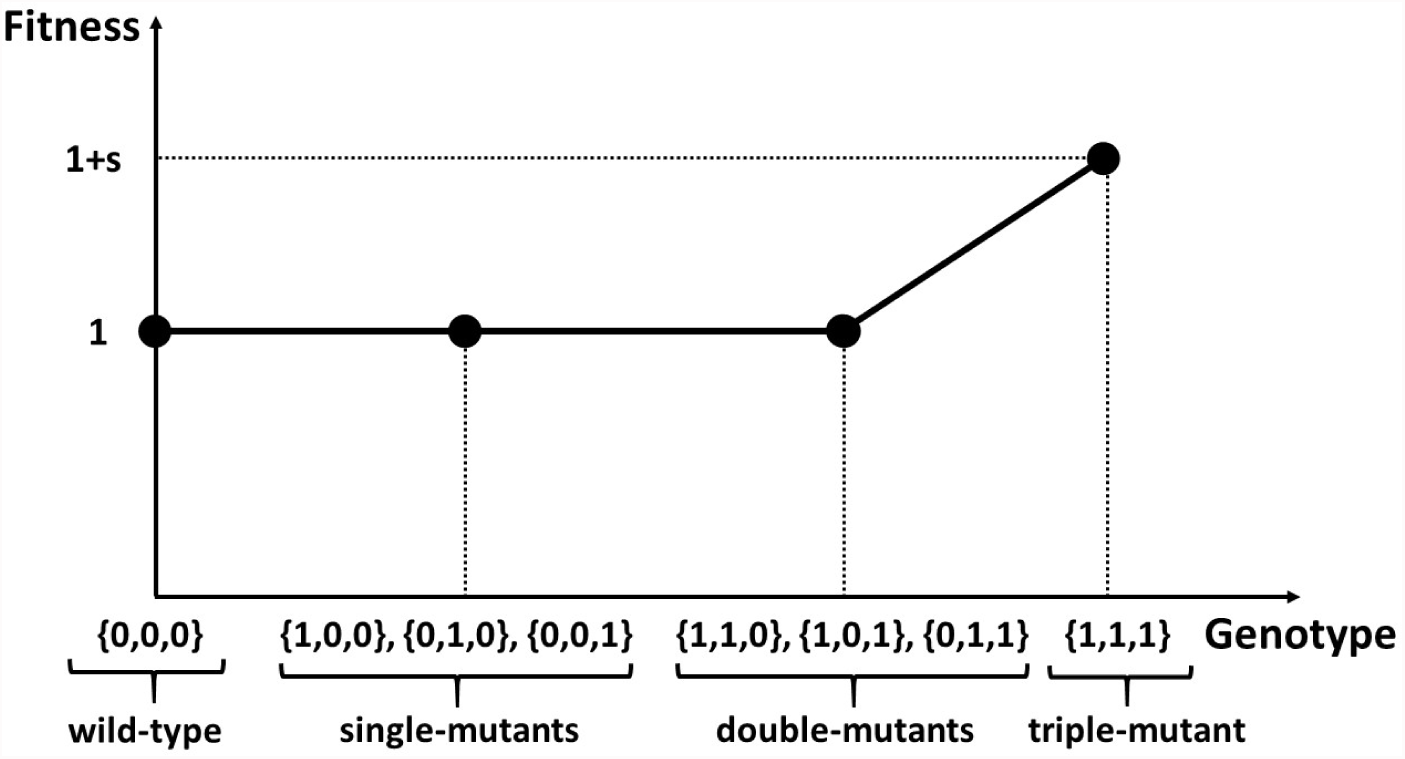
A visualization of the fitness landscape in the case of *K* = 3. The nodes represent the fitness of wild type, single-, double-, and triple-mutant genotypes. Wild-type alleles are denoted by 0 and mutants by 1. The {1,1,1} genotype has a fitness 1 + *s* > 1 and the rest have fitness 1.

**Table 1:** Symbol definitions

| Symbol | Definition |
| --- | --- |
| $N$ | Haploid population size |
| $\mu$ | Mutation rate per locus per generation |
| $r$ | Recombination rate per generation |
| $s$ | Selective advantage of the all-mutant genotype |
| $\mathcal{T}$ | Expected time until the all-mutant genotype makes up the majority of the population |
| $R_i(t)$ | The rate at which $i$-mutant individuals arise in the population at time $t$ |
| $p_i(t)$ | The probability that an $i$-mutant lineage arising at time $t$ will be successful |

## 3 Results

There are two fundamentally different plateau-crossing dynamics, depending on the relative rates of selection and recombination Eshel and Feldman [1970], Feldman [1971], Suzuki [1997], Jain [2009]. If recombination is weak relative to selection (*r* ≪ *s*), the adaptive genotype is rarely broken up by recombination and can spread rapidly once formed even if the individual mutant alleles are very rare in the population. If, on the other hand, recombination is strong (*r* ≫ *s*), the population is kept in quasi-linkage equilibrium, with the dynamics determined by the allele frequencies. Because the dynamics are so different, we consider these two regimes separately. Figure 2 and Table 2 summarize the different possible scaling behaviors of 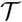 over all of parameter space.

**Figure 2:**
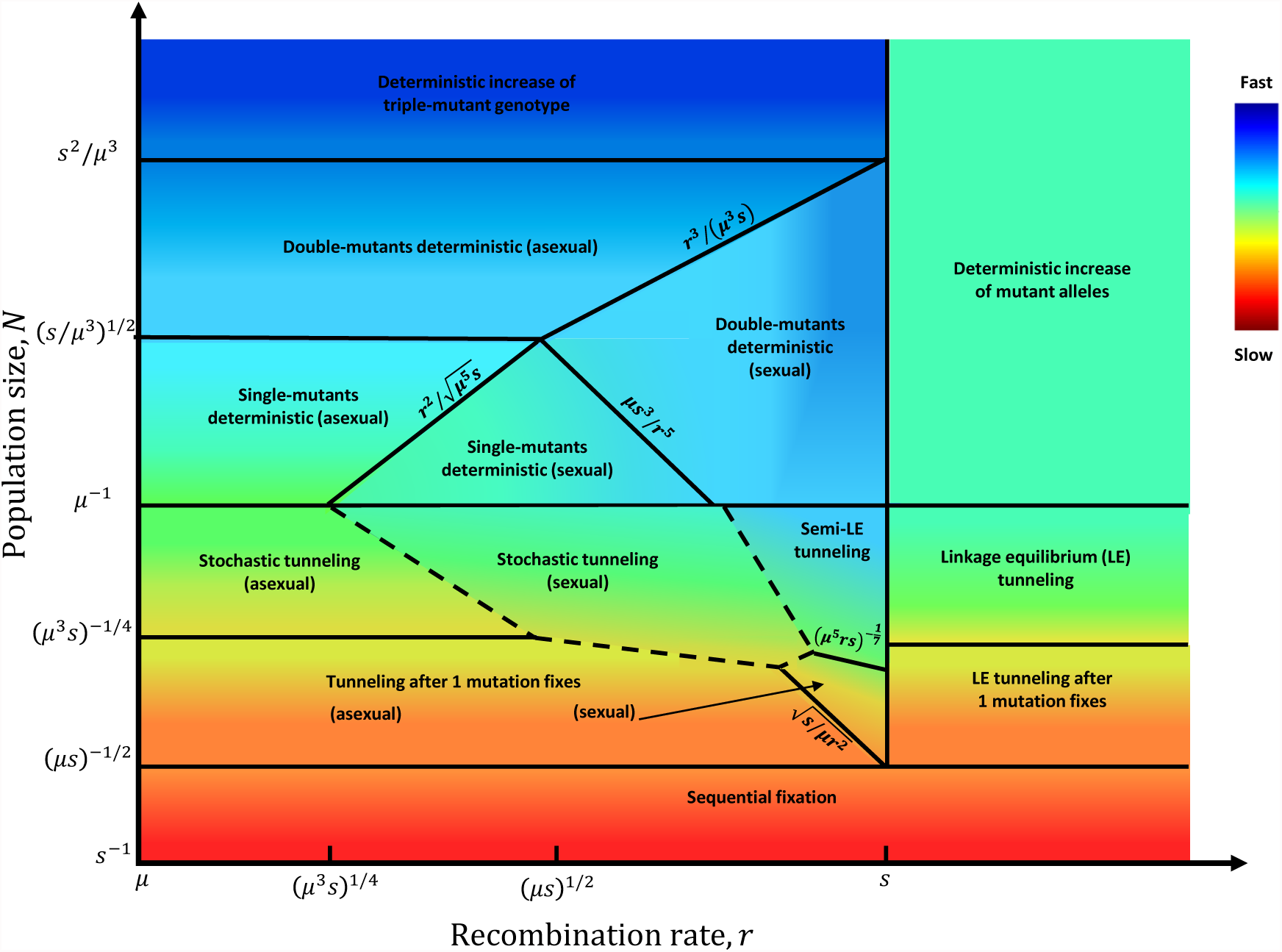
Schematic diagram of the asymptotic dynamics by which a population crosses a three-mutation (*K* = 3) fitness plateau to acquire a complex adaptation providing advantage *s*, as a function of recombination rate *r* and population size *N.* Color qualitatively represents the expected time 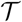 for the population to cross; for quantitative expressions, see Table 2. For *r* ≪ *s*, selection can drive the triple-mutant genotype to fixation while the other mutant genotypes remain rare, while for *r* ≫ *s* the population always remains close to linkage equilibrium; plateau crossing is fastest for intermediate recombination rates. The time to cross the plateau decreases as population size increases from the “sequential fixation” regime to the “deterministic” regimes. The “stochastic tunneling (sexual)” regime is a combination of several different regimes that can be practically indistinguishable, with boundaries that depend on the value of *μ*/*s* – see Appendix A.1.

**Table 2:**
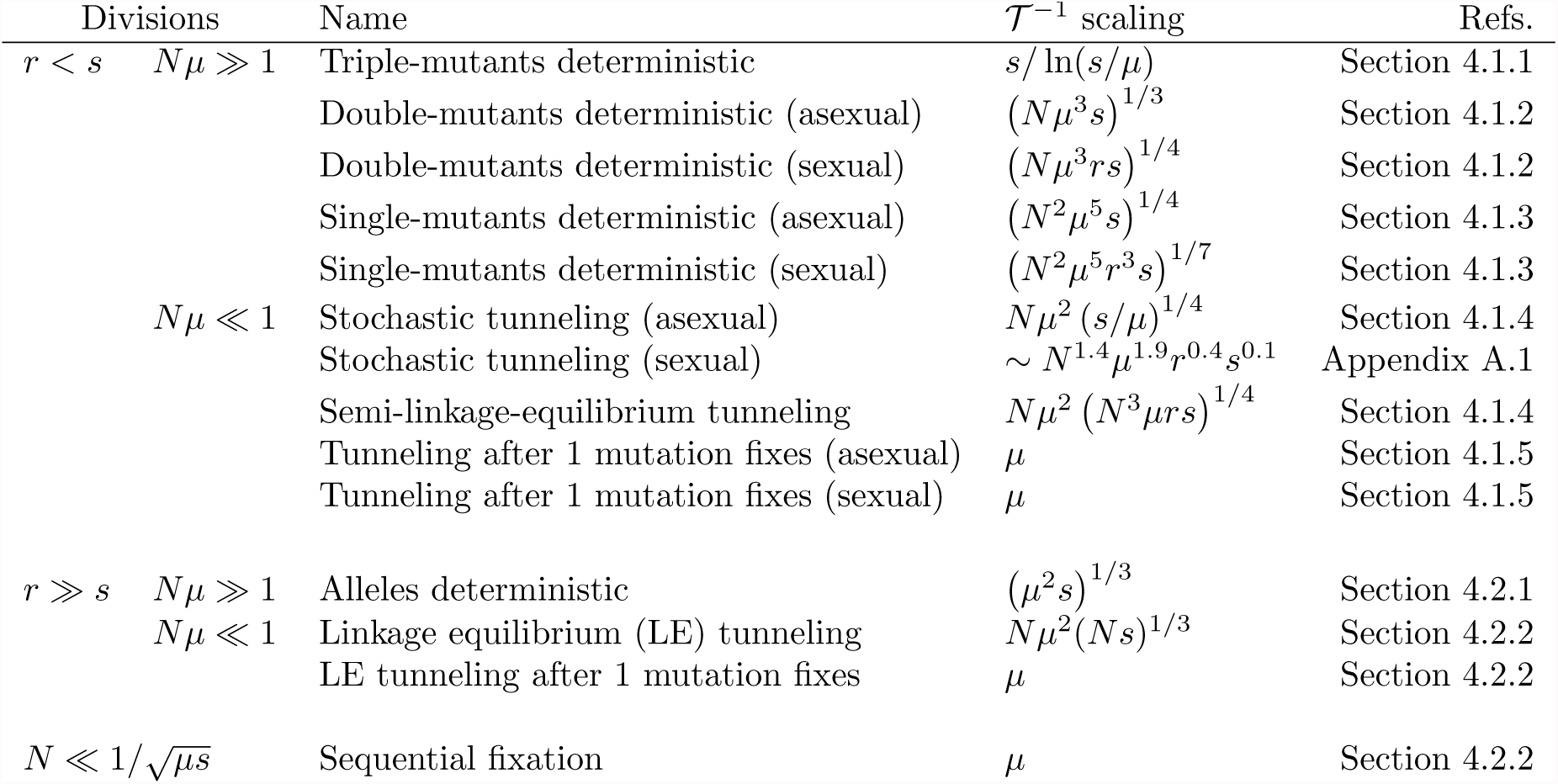
Rate of plateau-crossing (inverse of expected crossing time 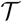) for asymptotic regimes shown in Figure 2, organized by main divisions of parameter space.

### 3.1 Rare recombination, *r* ≪ *s*

For *r* ≪ *s*, we must track genotype frequencies rather than just allele frequencies, so the complexity of the dynamics increases rapidly with the width of the plateau; we therefore focus on the simplest case that has not yet been fully characterized, *K* = 3. Even for this simplest case, there are many possible dynamical regimes (left half of Figure 2, Figure 3), depending on how difficult it is for the population to generate the adaptive genotype. For all but the largest population sizes, plateau-crossing becomes faster with increasing recombination rates, so the optimal rate is *r* ≲ *s.* The equations in this section all apply to this regime as well, with *s* replaced by the average rate of increase of triple-mutants when rare, 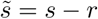.

**Figure 3:**
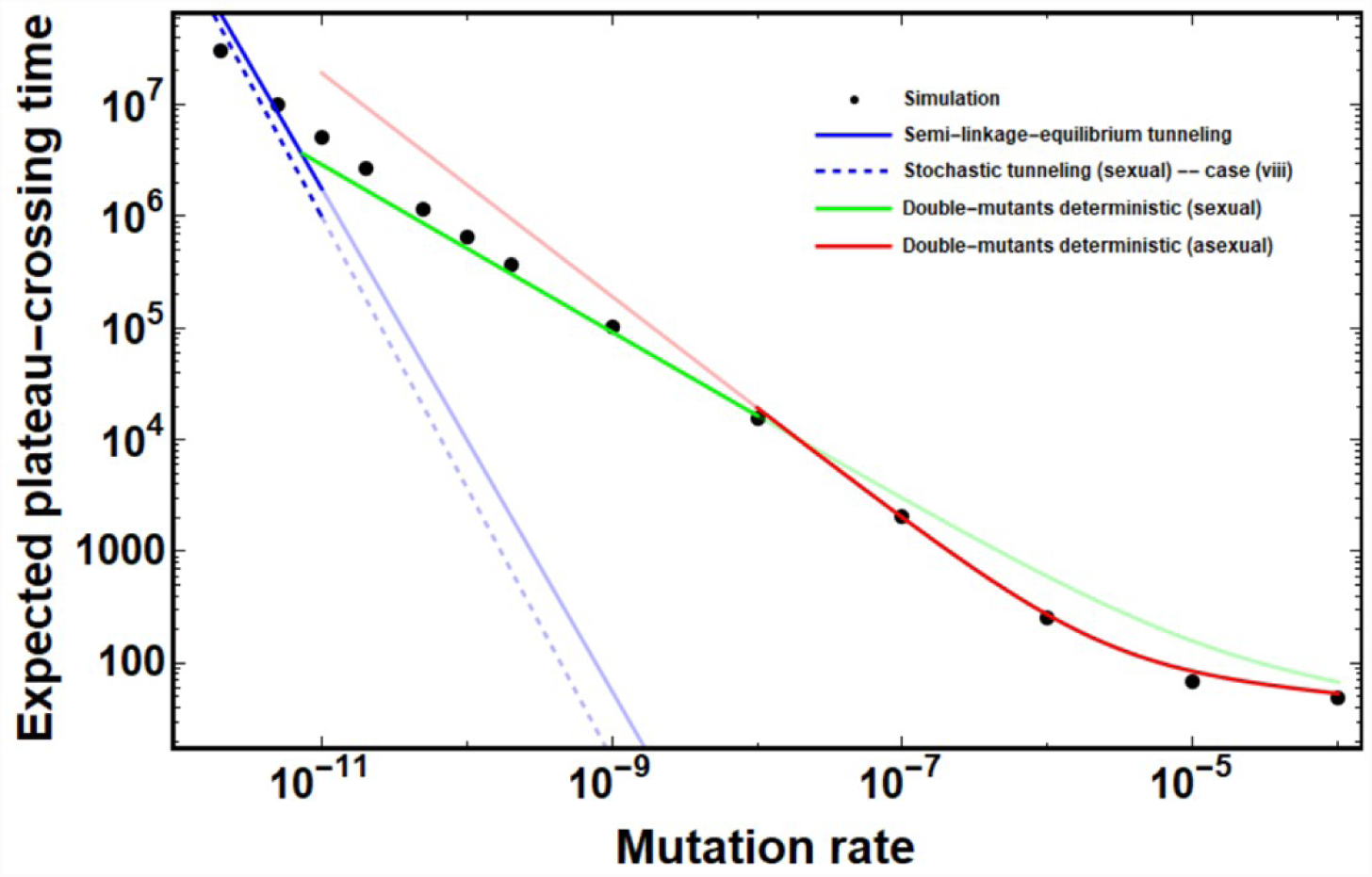
Expected time 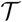 for a rarely recombining population to cross a three-mutation fitness plateau, as a function of the mutation rate *µ*. Points show simulation results, curves show analytical predictions. Note that in some regions, several analytical expressions give almost the same exact expected time. Typical plateau-crossing dynamics of the different asymptotic regimes are illustrated in Figures 5 and 6. The analytical solution for the double-mutants deterministic asexual and sexual paths (red and green), semi-linkage-equilibrium tunneling (solid blue) and single-mutant stochastic tunneling (blue dashed line) regimes is given by Equations 6, 11, and 27, respectively. Parameter values: *N* = 10^11^, *r* = 10^−3^, *s* = 1. Error bars are smaller than the size of the points.

If mutation and recombination are so frequent and the population is big enough that the triple-mutant genotype is generated effectively instantaneously, then the expected plateau-crossing time 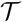 is just the time for a selective sweep, and depends primarily on *s.* For smaller *μ, r*, and *N*, most of 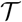 is waiting for the successful triple-mutant to be produced and the strongest dependence is on *μ. N* and *r* are most important at intermediate levels of diversity, where producing triple-mutants is difficult but there are opportunities for simultaneous polymorphisms at multiple loci to recombine. Quantitatively, when the mutation supply is large (*N_μ_* ≫ 1), then the expected plateau-crossing time is approximately:
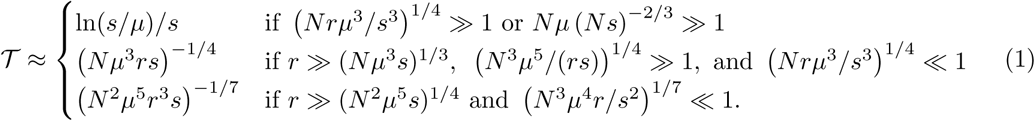

The first line of Equation 1 corresponds to the approximately deterministic dynamics of very large populations, which are insensitive to rare recombination (because the only substantial linkage disequilibrium is that generated by selection on the triple mutants after they are already on their way to fixation). In the second line, fluctuations in the number of triple-mutants are important, but single-and double-mutants can be treated deterministically (the “doubles deterministic (sexual)” regime). In the third line, fluctuations in the numbers of both triple- and double-mutants are important, but single-mutants can still be treated deterministically (the “single deterministic (sexual)” regime). If the recombination rate is lower than the thresholds in the second and third lines of Equation 1, the population is effectively asexual and 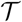 follows the scaling behavior described in Weissman et al. [2009] and Equations 6 and 9 below.

If the mutation supply is low (*Nμ* ≪ 1), then 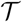 is approximately the expected waiting time for the first successful mutation. Since exploration of genotype space is more of a challenge for populations when mutations are rare, recombination has the potential to make more of a difference. When the recombination is very rare, the population is effectively asexual, with plateau-crossing rate 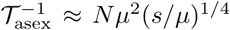 (Equation 10, see also Weissman et al. [2009]). As the recombination rate increases, it becomes easier mutations to be successful, and plateau-crossing speeds up. There are eight different asymptotic scaling regimes for rare recombination as *N* → ∞, depending on exactly how *μ, r, s* → 0, but for reasonable parameter values they are generally fairly similar (see Appendix A.1), with the expected rate of plateau-crossing roughly given by 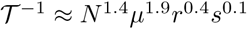. As recombination becomes more frequent (but still *r* ≪ s), pairs of large single-mutant lineages are able to succeed by reaching linkage equilibrium with each other and then recombining with a smaller third lineage (“semi-linkage-equilibrium tunneling”), and the rate of crossing increases further to 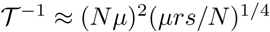 (Equation 11). This is the regime where recombination speeds plateau-crossing the most; comparing Equations 10 and 11, we see that it increases the rate by a factor ~ (*N*^3^*μ*^2^*r*)^1/4^, which could exceed an order of magnitude if *Nr* > 10^6^.

### 3.2 Frequent recombination, *r* ≫ *s*

For frequently recombining populations (*r* ≫ *s*), we find the expected time for plateau crossing across the full spectrum of possible plateau widths *K*, mutation rates *μ*, population sizes *N*, and selective coefficients *s* (Figure 4). These population will be in quasi-linkage equilibrium and selection will therefore act on alleles rather than genotypes. In this regime, the plateau-crossing time depends primarily on the mutation rate and is typically 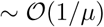. When mutations are frequent (*Nμ* ≫ 1), the population crosses the plateau nearly deterministically and solving the deterministic mutation-selection dynamics gives plateau-crossing time: 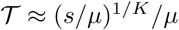.

**Figure 4:**
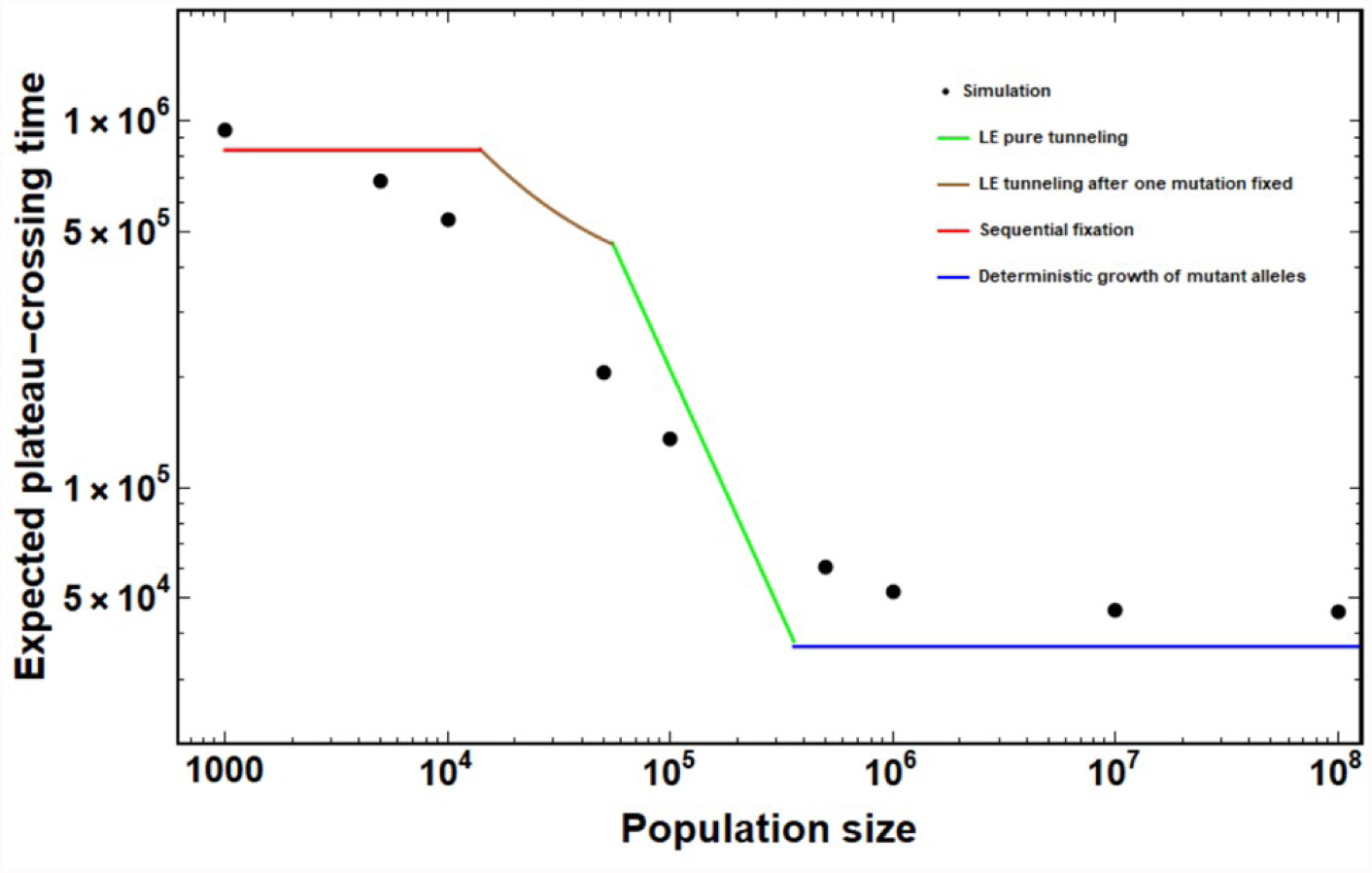
Expected time 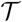 for a frequently recombining population to cross a three-mutation fitness plateau, as a function of the population size *N.* Points show simulation results, curves show analytical predictions Equation 13 (blue) and Equation 18 (green, brown, and red). Parameter values: *r* = 0.5, *μ* = 10^−6^, *s* = 0.05. The time to cross the plateau depends strongly on *N* for *N* ≲ 1/*μ*, and levels off for large or very small populations. Error bars are smaller than the size of the points.

**Figure 5:**
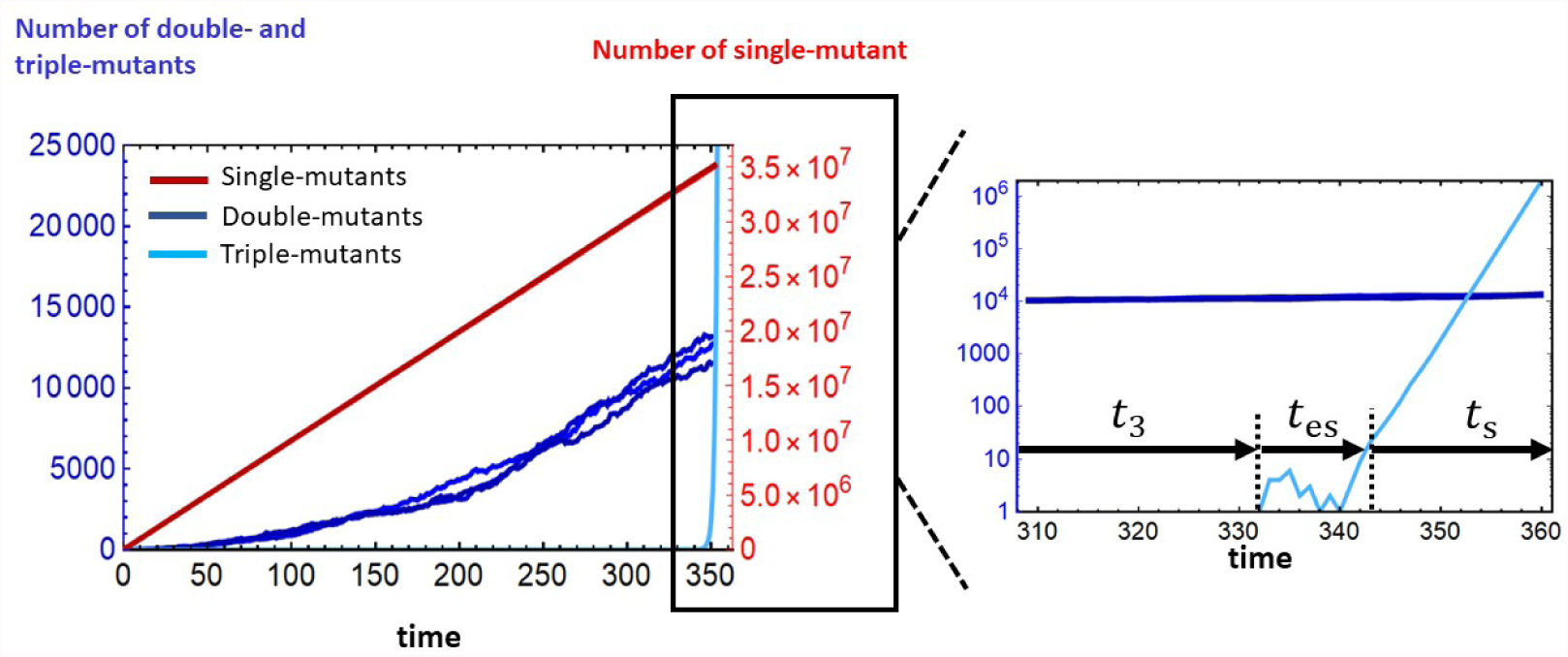
Typical simulation results of plateau-crossing dynamics for very large population sizes where 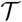 is dominated by the waiting time for the arrival of the first successful triple-mutant individual that has been produced following a mutation event in the growing population of double-mutants. The inset is a magnified view of the last 50 generations before the adaptive genotype fixes in the population which demonstrates the establishment time and sweep time of the triple-mutants. The model parameters for this simulation are *N* = 10^11^, *μ* = 10^−6^, *r* = 10^−3^, and *s* = 1.

**Figure 6:**
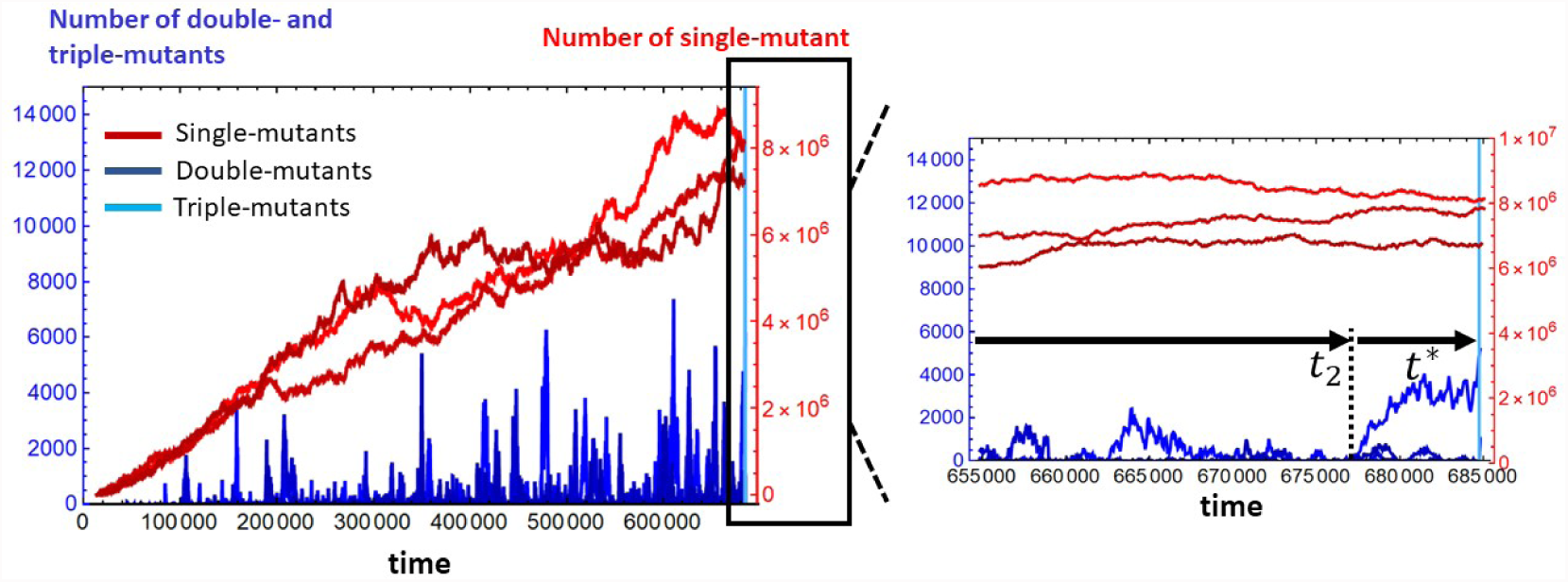
Typical simulation results of plateau-crossing dynamics for moderately large population sizes where 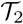 is dominated by the waiting time for the arrival of the first successful double-mutant lineage *t*_2_ (indicated on the inset) that has been produced following a recombination event between two single-mutants. The inset is a magnified view of the last few thousands of generations before the adaptive genotype fixes in the population which demonstrates that while the successful double-mutant lineage is drifting for *t*\* generations, the population of single-mutants has, to a very good approximation, remained constant. The model parameters for this simulation are *N* = 10^11^, *μ* = 5 × 10^−10^, *r* = 10^−3^, and *s* = 1.

When mutations are rare (*Nμ* ≪ 1), stochasticity is important and the dynamics typically proceed in two stages: first, *K* − *m* of the necessary mutations sequentially drift to fixation by chance; then, once the population is sufficiently close to the adaptive genotype, it relatively rapidly acquires the last *m* mutations together via stochastic tunneling. The typical value of *m* is the largest integer such that the probability of a new mutation triggering a tunneling event of *m* mutations is higher than the probability 1/*N* of fixation by drift:
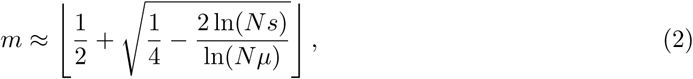

where ⌊.⌋ represents the floor function. Therefore, the plateau-crossing time is typically dominated by the time for *K* − *m* mutations to drift to fixation, unless *m* ≥ *K*, in which case the population tunnels directly. Summarizing these regimes, the expected time for a frequently recombining population to cross a fitness plateau is:
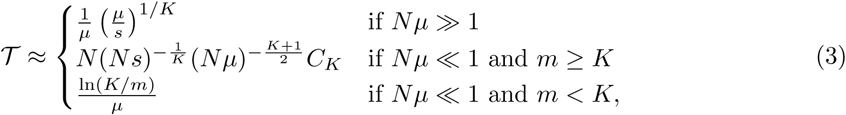

where *C_K_* in the second line (pure tunneling) is a combinatorial factor that depends only on *K* (see Equation 16).

Comparing the Equation 3 to the expected time for an asexual population to cross the (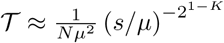 for *N* ≪ 1/*μ*, 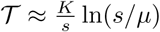 for *N* ≫ *s*^*K*−1^/*μ*^*K*^, with additional asymptotic regimes for intermediate population sizes [Weissman et al., 2009]), we see that frequent recombination tends to speed up adaptation in small populations (relative to asexuality), where the primary challenge is producing the beneficial genotype, while slowing it down in large populations, where most of the time is spent fixing the genotype after it has been produced.

## 4 Analysis

### 4.1 Rare recombination (*r* ≪ *s*)

In this section we will consider the plateau-crossing process in populations with rare recombination, starting with very large populations and progressively decreasing in size. As *N* decreases, the population’s ability to efficiently explore genotype space (measured by *N, μ*, and *r*) becomes more important, and its ability to exploit its discoveries (*s*) less so. At the largest population sizes, 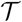 is essentially determined by *s.* For all the lower population size regimes, there will be at least some genotypes that are only rarely produced, and 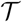 will be approximately the waiting time for the production of the first successful lineage of a rare genotype.

#### 4.1.1 Very large populations: deterministic dynamics

For extremely large population sizes, the number of single-, double-, and triple-mutant individuals are well approximated by their expected values after only a few generations. Triple mutants are produced almost instantaneously, and the plateau-crossing time is dominated by the time it takes them to sweep to fixation. This can easily be found by solving the deterministic equations for the dynamics of the genotype frequencies under mutation and selection, with recombination only reducing the effective selective advantage of the triple mutants:
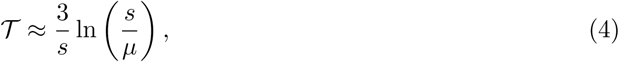

It is straightforward to generalize Equation 4 to arbitrary plateau widths *K:*

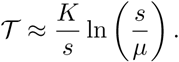

For this deterministic approximation to be accurate, the production rate of triple mutants, *R*_3_(*t*), must be large at the time *t* ~ 1/*s* when the first triple-mutant lineage reaches number 1/*s* and becomes established. Triple mutants are produced by double-mutant individuals that either acquire another mutation or recombine with single mutants, so *R*_3_ is:
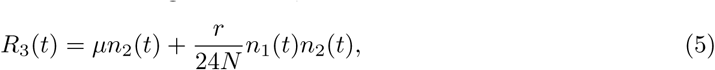

where *n_i_* is the total number of *i*-mutant individuals, so *n*_1_(*t*) ≈ 3*Nμt* and *n*_2_(*t*) ≈ 3*Nμ*^2^*t*^2^. (The factor of 1/24 in Equation 5 is because to make a triple mutant, each double mutant can only successfully recombine with 1/3 of the single mutants, and only 1/8 of the offspring will inherit the correct alleles.) At *t* ~ 1/*s*, Equation 5 gives *R*_3_ ≈ 3*Nμ*^3^/*s*^2^. (The recombination term is smaller by a factor 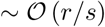 So to be in this regime, the population must have size *N* ≫ *s*^2^/*μ*^3^. Note that recombination is almost irrelevant in this regime: mutants are produced so frequently that there is no need for recombination to generate new combinations. (It does slightly slow down adaptation, as technically *s* should be replaced in the equations by 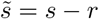.)

#### 4.1.2 Large populations: single- and double-mutants common, triple-mutants rare

Slightly smaller populations (with *N* ≪ *s*^2^/*μ*^3^) will still only occasionally be producing triple-mutants at the time they cross the plateau, so while the single- and double-mutant populations will have nearly deterministic dynamics, fluctuations in the number of triple-mutants will be important. Because triple-mutant lineages are rare, we can consider them in isolation, and 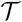 will be the waiting time for the first successful triple-mutant. The probability that a successful triple-mutant lineage will have been produced by time *t* is 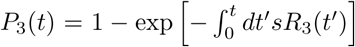, using that the probability that a triple-mutant lineage is successful once it has been produced is ~ *s* Ewens [2004]. Therefore, the expected waiting time 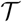 is given by:
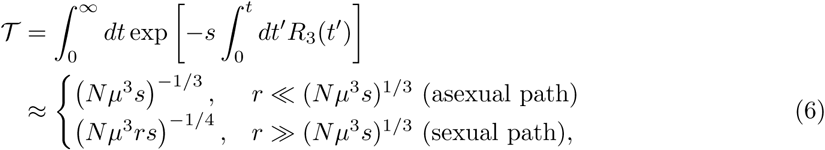

where we are ignoring constants of 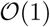. In the first line, the population is effectively asexual, i.e., the successful triple-mutant is likely to arise via mutation from a double-mutant. In the second line, it is more likely to arise via recombination between a double-mutant and a single-mutant. The 1 two expressions in Equation 6 are generally close, differing by a factor of only 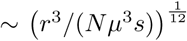: recombination can provide a mild increase in speed, but as in the previous section, the population is so large that the triple-mutant genotype will rapidly be produced by mutation alone anyway.

In deriving Equation 6, we ignored fluctuations in the number of double-mutants (as well as of single-mutants), so these must be negligible on time scales similar to 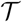 for the expressions to be valid. There are two ways that this approximation can hold. First, if the number of double-mutants is in fact close to its expectation with high probability. This will be true if the production rate *R*_2_ (*t*) of double-mutants is high, i.e., 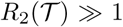, so that the double-mutant population is composed of many lineages and fluctuations in the individual lineage sizes average out. *R*_2_ is given by:
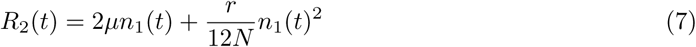

Plugging in the first line of Equation 6 for *t* gives the requirement 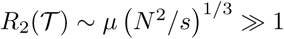

In the recombination-dominated regime in the second line of Equation 6, there is an additional way for the fluctuations to be negligible: recombination can cap their size by preventing them from greatly exceeding linkage equilibrium with the much larger and approximately deterministic wild-type and single-mutant populations. In this case, if *R*_2_ ≪ 1 the number of double-mutants will be fluctuating as lineages are sporadically produced and die out, but no one lineage will drift for a time much exceeding ~ 1/*r* before being broken up by recombination. We will see what condition this puts on the parameters in the following section, but for now, note that this mechanism requires the single-mutants to be approximately deterministic, so at a minimum we require *R*_1_ ~ *Nμ* ≫ 1.

Finally, we must also check the conditions for our assumption that the triple-mutant lineages are rare enough to be considered in isolation. This is equivalent to 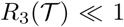 – the flip side to the parameter condition in the preceding section requiring that triple-mutants be approximately deterministic. Plugging in our expressions for *R*_3_ and 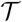, we get *μ*(*N*/*s*^2^)^1/3^ ≪ 1, which is indeed the reverse of the previous condition.

#### 4.1.3 Moderately large populations: single-mutants common, double-mutants rare

For populations slightly smaller than those in the previous section, the mutation supply will still be high (*Nμ* ≫ 1), so single-mutants will still be approximately deterministic, but double-mutant lineages will be rare 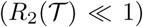 and we must consider their fluctuations. Since they are rare, we can consider each lineage in isolation, and 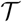 will be the waiting time for the first successful double-mutant to arise.

A double-mutant lineage can be successful by either mutating or recombining with single-mutants to produce a successful triple-mutant. Since the single-mutants are deterministic (*n*_1_(*t*) ≈ 3*Nμt*), we can lump these two processes into a single time-dependent effective mutation rate, 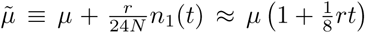. Since in this regime we expect the waiting time for the first successful double-mutant lineage to be long compared to the time for which that lineage must drift before producing the successful triple-mutant, we can further treat this effective mutation rate as being approximately constant over each lineage’s lifetime. With this approximation, the problem is reduced to that considered in Weissman et al. [2009]: a lineage mutates at rate 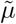 to a genotype with advantage *s*; additionally, the double-mutant lineage has an effective selective disadvantage *r* due to being broken up by recombination with the wild type. The probability *p*_2_(*t*) that a double-mutant lineage arising at time *t* will be successful is therefore [Weissman et al., 2009]:
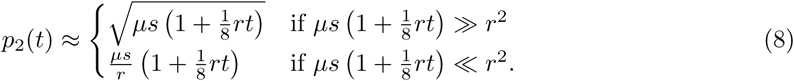

In the first line of Equation 8, a lineage is most likely to succeed by drifting for long enough to produce many (~ 1/*s*) triple-mutants. In the second line, recombination is too frequent and lineages are broken up before they can drift for that long. We therefore see that the condition for being able to ignore fluctuations in the double-mutant numbers as in the previous section 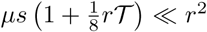. Since this case is covered by that section’s analysis, we will now focus on the case 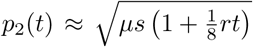 where fluctuations are key. Combining the success probability *p*_2_(*t*) with the production rate 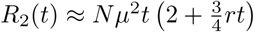, we can find the expected waiting time 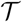 for the first successful double-mutant (ignoring 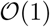 constants):
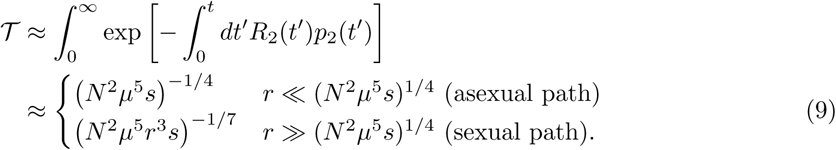

Both *R*_2_ and *p*_2_ switch from being mutation-dominated to recombination-dominated at time *t* ~ 1/*r.* In the first line of Equation 9, 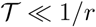 so the population is effectively asexual. In the second line, 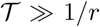 so the successful double-mutant is likely both to be produced by recombination and to produce the successful triple via recombination.

#### 4.1.4 Moderately small populations: occasional triple polymorphisms

If the mutation supply is low (*Nμ* ≪ 1), then the population will typically be monomorphic. The plateau-crossing time is dominated by the waiting time for a lucky single-mutant lineage that drifts long enough to either fix or encounter additional mutations that allow it to tunnel across the plateau. We will consider the latter process in this section. Call this mutation *A*, and let *T_A_* be the time for which this mutation’s lineage must drift to be likely to be successful; over this time, the lineage will typically reach a size *n_A_* ~ *T_A_.* The mutation will manage to drift this long with probability *p*_1_ ~ 1/*T_A_*, so the expected plateau-crossing time is 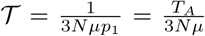. Note that if *T_A_* > *N* (or, equivalently, 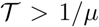), the lineage is more likely to fix than tunnel. We now find expressions for the necessary drift time *T_A_.*

First, we will review the asexual process. Weissman et al. [2009] showed that *T_A_* ~ (*μ*^3^*s*)^−1/4^ (ignoring combinatoric factors) is long enough for the lineage to be likely to acquire two additional mutations (which we will call *B* and *C*) and be successful. The expected time to cross the plateau is thus:
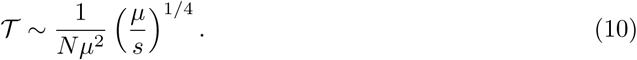

Comparing *T_A_* to *N*, we see that the population will only tunnel if *N* (*μ*^3^*s*)^−1/4^ > 1. This result therefore applies only to populations within a fairly narrow band of sizes, with the lower limit of validity only a factor of (*μs*)^1/4^ smaller than the upper limit – less than three orders of magnitude for realistic parameters.

Recombination can speed up tunneling (i.e., reduce the necessary *T_A_*) by allowing the original *A* lineage to acquire *B* and *C* from the ~ *NμT_A_* independent mutant lineages that arise while it is drifting. Let *B* be the mutation carried by the largest such lineage; it will typically drift for *T_B_* ~ *NμT_A_* generations, reaching a size *n_B_* ~ *NμT_A_.* If *T_B_* ≫ 1/*r*, recombination will effectively reduce the linkage disequilibrium between *A* and *B*, i.e., there will be an average of 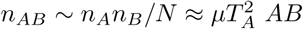 individuals over most of the *T_B_* generations for which both mutations are drifting. During this time, there will be ~ *NμT_B_ C* lineages produced by mutation, the largest of which will therefore typically drift for *T_C_* ~ *NμT_B_* ≈ (*Nμ*)^2^*T_A_* generations to size *n_C_* ~ *T_C_. AB* and *C* individuals will therefore coexist for ~ *T_C_* generations, during which they will generate 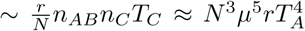 triple-mutant recombinants. Each of these has a probability ≈ *s* of being successful, so we see that for our original *A* lineage to be likely to be successful, its drift time *T_A_* must satisfy 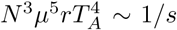. Solving for *T_A_* gives *T_A_* ~ (*N*^3^*μ*^5^*rs*)^−1/4^, corresponding to an expected plateau-crossing time of:
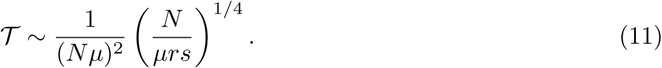

We refer to this as “semi-linkage-equilibrium tunneling”, since the two most frequent mutations are in linkage equilibrium with each other while drifting, but the third mutation may not be, and the triple-mutant will produce large linkage disequilibria once it starts to sweep.

The derivation of Equation 11 assumed that *A* and *B* were close to being in linkage equilibrium with each other, i.e., *rT_B_* ≫ 1. Substituting in *T_B_* ~ *NμT_A_*, this is equivalent to a condition that *T_A_* ≫ 1/(*Nμr*). However, it may be the case that the *A* and *B* lineages can produce enough recombinants to be successful before they approach linkage equilibrium. This is true for small values of *r*, where the time to approach linkage equilibrium becomes very long. In this situation, the analysis here overestimates how large *T_A_* must be, and therefore overestimates the time required to cross the plateau. The correct analysis of this regime is even more involved, and we leave it for Appendix A.1. We also ignored the possibility that the *AB* individuals might produce a triple mutant directly via mutation, but it is straightforward to check that this is rare in the relevant parameter range: as long as 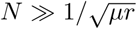, acquiring the third mutation via recombination is more likely.

#### 4.1.5 Small populations: single-mutants drift to fixation

For smaller populations, the most likely way for a single-mutant to be successful is for it to drift to fixation, which occurs with probability 1/*N.* The expected waiting time is therefore 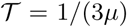. Once the single-mutant has fixed, the population only needs two additional mutations, so Weissman et al. [2010]’s two-locus analysis applies. The average time to tunnel will necessarily be small compared to the time for the first mutant to drift to fixation, so it can be neglected in 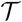. The exception is for very small populations, 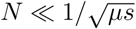, where the *second* mutation is also more likely to drift to fixation than to tunnel [Weissman et al., 2010]. In this case, the total waiting time is 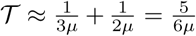. (The final fixation of the third mutation is relatively rapid as long as *Ns* ≫ 1.)

### 4.2 Frequent recombination (*r* ≫ *s*)

If recombination is frequent (*r* ≫ *s*), selection will be too weak to generate linkage disequilibrium, and the population will stay close to linkage equilibrium (LE). We can therefore simply track allele frequencies, rather than genotype frequencies. For this much easier problem, we can consider plateaus of arbitrary width *K.*

#### 4.2.1 Large populations (*Nμ* ≫ 1): deterministic dynamics

When the mutation supply is large, *Nμ* ≫ 1, the mutant allele frequency trajectories are nearly deterministic, and therefore almost the same as each other, i.e., a single variable *x*(*t*) can describe the frequency of all the mutant alleles. When the mutations are rare (*x* ≪ 1), their increase according to:
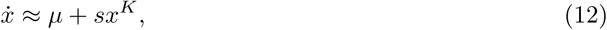

where *x^K^* is the frequency of the beneficial genotype. Solving Equation 12 for *t* such that *x*(*t*) ≈ 1 gives the time to cross the plateau:
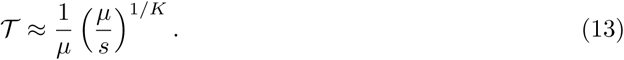

We can understand this as the time it takes for mutation to drive the mutations to the frequency *x* ~ (*μ*/*s*)^1/*K*^ at which selection takes over, after which fixation is rapid.

#### 4.2.2 Small populations (*Nμ* ≪ 1): sequential fixation + stochastic tunneling of mutant alleles

When the mutational supply of the population is small (*Nμ* ≪ 1), most loci will usually be monomorphic, with occasional drifting mutant lineages. To cross the plateau, the population needs some combination of mutations drifting to fixation, and others producing the beneficial genotype and tunneling together. We can think of the tunneling dynamics as allowing the population to “see” the adaptive genotype once the dominant genotype is within *m* mutations of it, for some *m.*

We must first find how the maximum tunneling range *m* depends on *N, μ*, and *s.* As in section **??**, a population can cross the plateau via a rare mutant lineage that grows to a large size over an extend period of time. Suppose that such a lineage persists for ~ *T*_1_ generations, typically growing to size ~ *T*_1_. There will be ~ (*m* − 1)*NμT*_1_ mutations at other loci during this time, the largest of which will typically persist for *T*_2_ ~ (*m* − 1)*NμT*_1_ while the original allele is still drifting. During *T*_2_, the longest-drifting mutation at a third locus will typically persist for *T*_3_ ~ (*m* − 2)*NμT*_2_ = (*m* − 1) (*m* − 2) (*Nμ*)^2^*T*_1_, and so on, with the the *m^th^* mutation persisting for *T_m_* ~ (*K* − 1)!(*Nμ*)^*m*−1^*T*_1_. The frequency *x_m_* of the *m*-mutant genotype will peak at:

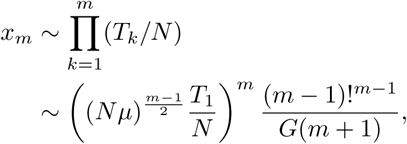

where *G* is the double gamma function. For the mutations to establish, this peak frequency must exceed ~ 1/*Ns.* Solving this condition for *T*_1_ gives the time scale over which the first mutation must drift to be likely to be successful:
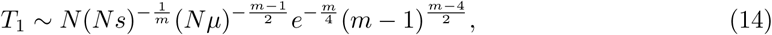

where the final combinatorial factors are approximations valid for large *m*, and negligible for small *m.* For the initial mutation to be more likely to tunnel than to fix, *T*_1_ must be small compared to *N.* Solving *T*_1_ ~ *N* for *m* therefore gives the maximum tunneling range:
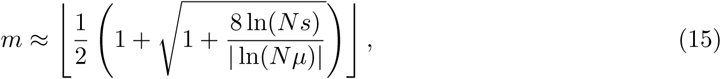

where ⌊.⌋ is the floor function.

If *m* ≥ *K*, then a wild-type population can tunnel directly to the beneficial genotype. The probability that a mutation successfully tunnels is ~ 1/*T*_1_, so the expected waiting time is:
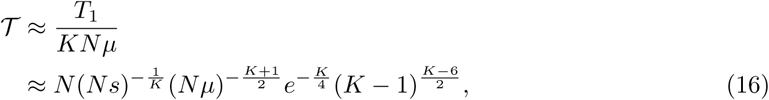

where we have substituted Equation 14 with *m* = *K* for *T_1_.* If 1 < *m* < *K*, the total plateau-crossing time is dominated by the time it takes for the population to fix *K* − *m* mutations via drift so that it can get close enough to the adaptive genotype to tunnel the rest of the way. (If *m* = 1, then the population cannot tunnel and must fix *K* − 1 mutations by drift, at which point the *K^th^* mutation becomes beneficial.) The *k^th^* mutation fixes after an expected waiting time of 1/(*K* − *k* + 1)*μ*, so the total expected waiting time for *K* − *m* mutations to fix is
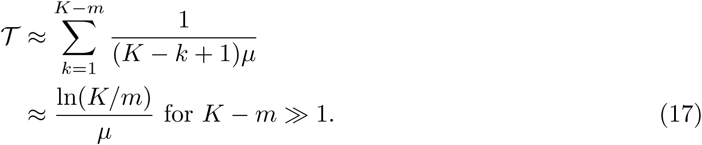

#### 4.2.3 Three-mutation plateaus

Plugging in *K* = 3 to the above analysis, the expected time to cross the plateau is (with 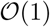 combinatorial factors included for clarity):
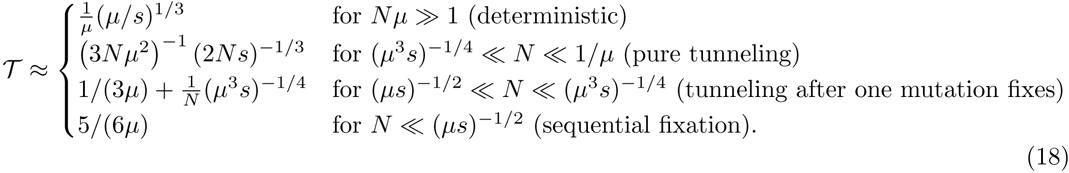

The second term in the third line is the *K* = 2 tunneling time [Weissman et al., 2010].

## 5 Discussion

We have shown that even moderately large populations can acquire complex adaptations requiring three individually-useless mutations substantially faster than would be expected if mutations had to fix sequentially by drift. In other words, natural selection can at least somewhat effectively promote mutations that not only provide no direct selective benefit, but also do not directly increase evolvability, i.e., do not change the distribution of mutational effects. Recombination helps most at intermediate population sizes, where there can be simultaneous polymorphisms at multiple loci but triple mutants are rare. In this range, the rate of plateau-crossing is maximized when recombination is just somewhat rarer than selection.

Across regimes, the rate of crossing the three-mutant fitness plateau scales sub-cubically in the mutation rate, i.e., complexity is not strongly suppressing the rate of adaptation, suggesting that even more complex adaptations could also potentially be acquired. However, analyzing even the three-mutation case for *r* ≲ *s* involved a proliferation of different dynamical regimes, so simply extending our analysis to wider plateaus is likely to be impractical. The asexual case Weissman et al. [2009] and the case *r* ≫ *s* analyzed above are simpler but plateau-crossing is fastest for *r* ≲ *s*, meaning that these easier limiting cases may be missing essential dynamics.

How practically important could adaptive paths across three-mutation plateaus be? Could we hope to observe experimental populations following them? Viruses often have large populations and high mutation rates; if we consider an RNA virus with a mutation rate of ~ 10^−4^ per base per replication and a potential adaptation providing a ~ 10% fitness advantage, a population size of *N* > 10^9^ – fewer than might be present in a single infected host – would be enough for the population to deterministically acquire the triple-mutant genotype. On the other hand, if we consider a yeast population in which the relevant mutations have target sizes of ~ 300 base pairs, for a mutation rate of ~ 10^−7^ Lynch et al. [2008], it would be difficult to maintain a large enough experimental population for long enough to reliably acquire the adaptation via any of the paths we have described here.

The major limitation in seriously applying any of our analysis to real populations is that we have considered the necessary loci in isolation. As mentioned in the Introduction, a major part of our motivation in considering the possibility of complex adaptation is that a combinatorial argument suggests that there are potentially very many of them available. But if there are in fact very many possible complex adaptations, then the first one that actually fixes in the population is likely to be one that happened unusually quickly, potentially by different dynamics than those considered here Weissman et al. [2010]. Thus at a minimum, we would need to consider the entire distribution of plateau-crossing times rather than just its mean. More precisely, we would need to describe the left tail of the distribution. This may in fact simplify the analysis – there may be only a few ways for a lineage to get a lot of mutations quickly, regardless of the population parameters Weissman et al. [2010] – and thus provide a way forward to analyzing wider plateaus.

The fact that the population is likely to be adapting at more than just *K* loci does not only mean that we need to think about the left tail of crossing-time distribution; it also means that we need to think about how adaptation elsewhere in the genome may affect evolution at the focal loci. If there is substantial fitness variance due to the rest of the genome and limited recombination, the dynamics of the mutant lineages will be completely different due to hitchhiking Neher and Shraiman [2011]. In addition, the complex adaptation may be lost due to clonal interference once it is produced, reducing its fixation probability. The fixation probability in this case is likely to require a careful calculation in its own right, as the background fitness is likely to systematically differ from the mean because of the required conditioning on long-lasting lineages carrying the intermediate mutations.

Perhaps even more importantly than clonal interference is the potential epistatic interference. When we consider just the *K* focal loci, substitutions at other loci should turn the fitness landscape into a constantly shifting “seascape” Mustonen and Lässig [2009]. In the most extreme case, other mutations may fix that permanently disrupt the potential complex adaptation, forcing the population onto another path. We have no understanding of how this should affect the probability of complex adaptation in anything beyond the simplest possible case of a single beneficial mutation blocking a two-mutation complex adaptation in an asexual population Ochs and Desai [2015]. We can already see that is likely to substantially change the interpretation of our results by looking at Table 2 and our results for generic *K* with *r* ≫ *s* and *Nμ* ≪ 1. The regimes in which the population only tunnels through the last mutations while initially fixing the others via drift appear to have roughly the same rate of plateau-crossing as the sequential fixation regime in which all mutations but the very last must drift to fixation. But this is because our model assumes that all populations reach the adaptive genotype eventually. In a more realistic model in which populations can get diverted and miss potential adaptations entirely, being able to tunnel through *m* mutations greatly increases the zones of attraction of adaptive genotypes in the fitness landscape, and could make a large difference in the probability of finding them.

In addition to epistatic interactions with other loci, the plateau could shift because of environmental changes Masel [2006], Kim [2007]. It is difficult to say which process is likely to be more important. We currently do not even know whether changes in the selective coefficient of single mutations are driven more often by environmental changes or changes in the rest of the genome, let alone what drives changes in selection on the rest of the genome. More generally, the basic difficulty in analyzing more complex, realistic fitness landscapes is that we have no idea what they should look like. Even mapping out the local fitness landscape of a single gene requires a heroic experimental effort (e.g., Bank et al. [2016]) – and then we only know it in a limited number of artificial environments. Our best hope may be to try to develop a theory that can reduce the full complexity of landscapes to a reasonable number of parameters describing their features that are most relevant for adaptation, but it is an open question whether such a theory exists.

## Acknowledgments

DW was supported in part by a Mathematical Modeling of Living Systems Investigator award from the Simons Foundation.

## A Appendix

### A.1 Small populations with rare recombination

Here we focus on populations with low mutation supply, *Nμ* ≪ 1, and rare recombination, *r* ≫ *s*. In particular, we focus on those that fall in between the asexual and semi-linkage equilibrium cases discussed above, for which recombination is frequent enough to speed plateau-crossing but too rare to bring even the largest mutant lineages into linkage equilibrium with each other. As above, the expected plateau-crossing time is dominated by the waiting time for the production of the first successful single-mutant lineage *A* which drifts for time *T_A_*, with the other possible mutations labeled *B* and *C.* All genotypes that drift for a time *T_X_* reach a typical size *n_X_* ~ *T_X_*, so we will not need to distinguish between drift times and lineage sizes in the following. We will exploit our freedom in labeling the *B* and *C* mutations to always label the double mutant *AB* if it has the *A* allele, so the *AC* genotype will not appear in our analysis. Throughout, we will ignore 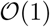 numerical factors arising from combinatorics and integration, none of which change the results significantly. We can identify eight possible asymptotic scenarios, depending on the relative sizes of the different relevant lineages (Figure 7):

**Figure 7:**
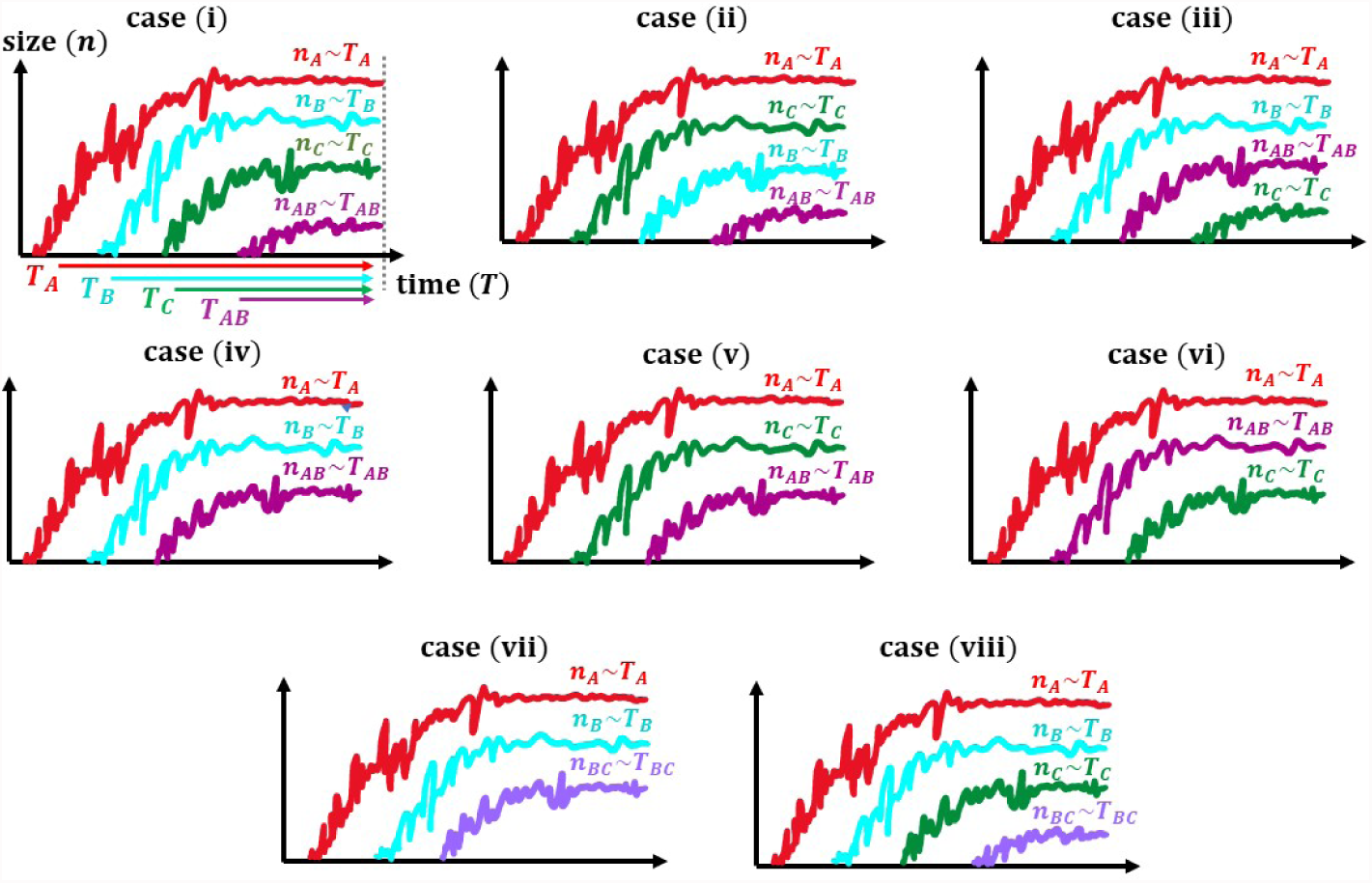
A schematic plot of 8 different ways in which the plateau-crossing can occur given that all the mutant alleles are not frequently produced in the population. In case (i), the two bigger lineages, *A* and *B*, recombine with each other to produce the *AB* lineage which in turns recombines with the *C* lineage to produce the first successful *ABC* lineage after *TA* generations. Cases (ii) and (iii) also correspond to the same dynamics as case (i) except for that the second and third biggest mutant lineages could be different. Case (iv) occurs when lineages *A* and *B* recombine to produce an *AB* lineage which later mutates to produce the beneficial mutant. In both cases (v) and (vi), the *A* lineage mutates to produce the *AB* lineage which then recombines with the *C* lineage to produce the successful *ABC* lineage. In case (vii), the *B* lineage mutates to produce a *BC* lineage which later recombines with *A.* Case (viii) occurs when the *B* and *C* lineages recombine to produce a *BC* lineage which then recombines with the *A* lineage.

(i) *T_A_* ≫ *T_B_* ≫ *T_C_* ≫ *T_AB_*: While three independent mutant lineages are drifting, the larger two recombine, and then that recombinant recombines with the third lineage to produce the successful triple-mutant. Typical sizes are *T_B_* ~ *NμT_A_, T_C_* ~ *NμT_B_*, and 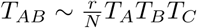 (because we only consider the largest *AB* lineage that arises while *C* is drifting). The number of *ABC* individuals produced by recombination between *C* and *AB* during the ~ *T_AB_* generations that they coexist is 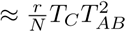; we need this quantity to be ~ 1/*s* for success to be likely:

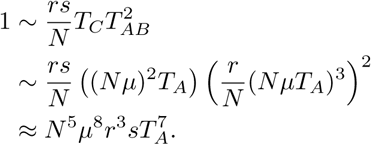

Solving for *T_A_* and the other drift times gives:
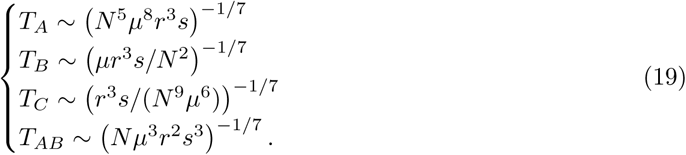

(ii) *T_A_* ≫ *T_C_* ≫ *T_B_* ≫ *T_AB_*: While three independent mutant lineages are drifting, the largest recombines with the smallest, and then that recombinant recombines with the middle lineage to produce the successful triple-mutant. Typical sizes are *T_C_* ~ *NμT_A_, T_B_* ~ *NμT_C_*, and 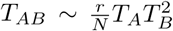. To get ~ 1/*s* triple-mutants, we need:

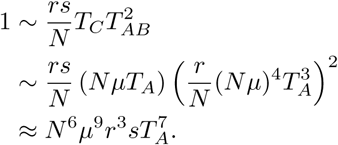

Solving for *T_A_* and the other drift times gives:
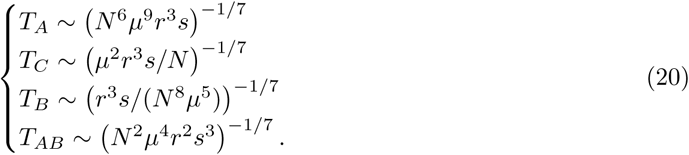

(iii) *T_A_* ≫ *T_B_* ≫ *T_AB_* ≫ *T_C_:* Two single-mutant lineages recombine. While that recombinant double-mutant is drifting, a third single-mutant lineages arises and recombines with it to produce a successful triple-mutant. Typical sizes are *T_B_* ~ *NμT_A_*, 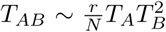, and *T_C_* ~ *NμT_AB_.* To get ~ 1/*s* triple-mutants, we need:

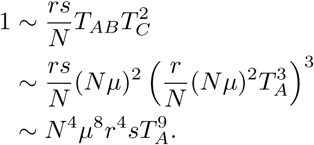

Solving for *T_A_* and the other drift times gives:
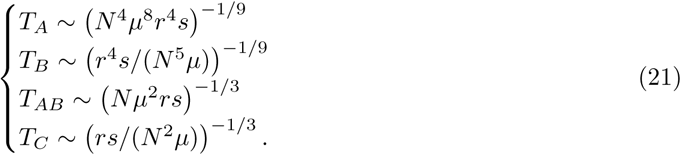

(iv) *T_A_* ≫ *T_B_* ≫ *T_AB_:* Two single-mutant lineages recombine, and that recombinant lineage then mutates and succeeds. Typical sizes are *T_B_* ~ *NμT_A_* and 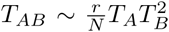. The *AB* lineage will produces 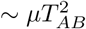 mutants while it is drifting; setting this equal to ~ 1/*s* gives:

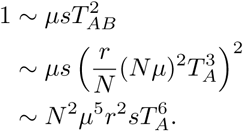

Solving for *T_A_* and the other drift times gives:
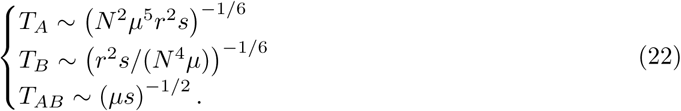

(v) *T_A_* ≫ *T_C_* ≫ *T_AB_*: While two single-mutant lineages are drifting, the mutates at the third locus. This double-mutant then recombines with the other single-mutant lineage. Typical sizes are *T_C_* ~ *NμT_A_* and *T_AB_* ~ *μT_A_T_C_* (because we only consider the largest-double mutant lineage that arises while *C* is drifting). To get ~ 1/*s* triple-mutants, we need:

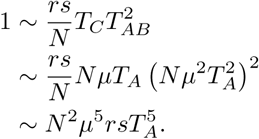

Solving for *T_A_* and the other drift times gives:
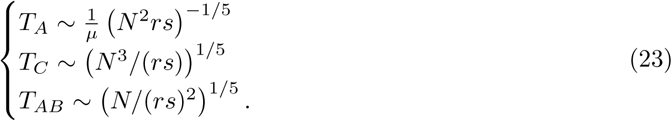

(vi) *T_A_* ≫ *T_AB_* ≫ *T_C_*: A single-mutant lineage mutates. While the resulting double-mutant lineage drifts, a new lineage with a mutation at the third locus arises and successfully recombines with it. Typical sizes are 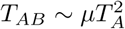 and *T_C_* ~ *NμT_AB_.* To get ~ 1/*s* triple-mutants, we need:

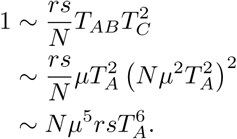

Solving for *T_A_* and the other drift times gives:
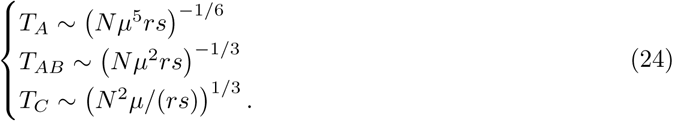

(vii) *T_A_* ≫ *T_B_* ≫ *T_BC_*: While two single-mutant lineages are drifting, the smaller one acquires an additional mutation at the third locus. This double-mutant lineage then successfully recombines with the larger single-mutant lineage. Typical sizes are *T_B_* ~ *NμT_A_* and 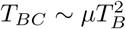. To get ~ 1/*s* triple-mutants, we need:

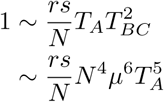

Solving for *T_A_* and the other drift times gives:
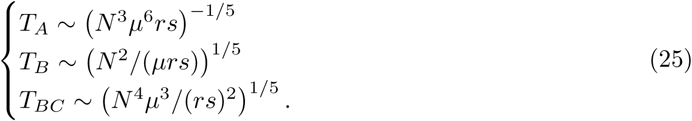

(viii) *T_A_* ≫ *T_B_* ≫ *T_C_* ≫ *T_BC_*: While three single-mutant lineages are drifting, the smaller two recombine. The recombinant then successfully recombines with the largest single-mutant lineage. Typical sizes are *T_B_* ~ *NμT_A_, T_C_* ~ *NμT_B_*, and 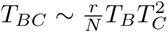. To get ~ 1/*s* triple-mutants, we need:

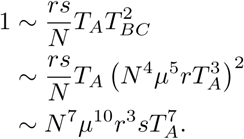

Solving for *T_A_* and the other drift times gives:
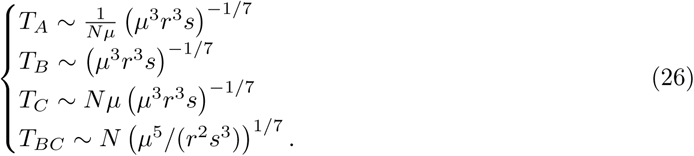

For all of these cases, the expected plateau-crossing time is 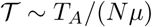. All require that the double-mutant drift times *T_AB_* or *T_BC_* be small compared to 1/*r*, so that the lineage is not broken up by recombination. We collect the predicted rates and conditions here:
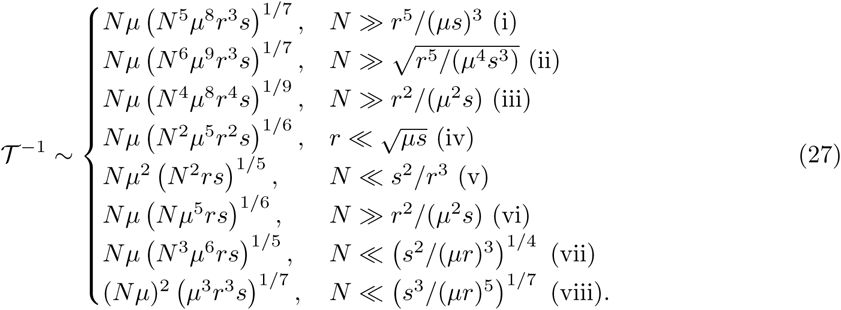

For parameter values where multiple cases apply, the predicted 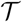 value is the one corresponding to the case with the smallest *T_A_* – the rates for the different cases do not add, since all are dependent on the same initial dynamic of an unusually long-lived single-mutant. If even the smallest *T_A_* is greater than *N*, single-mutants are more likely to fix than tunnel. For most reasonable parameter values, multiple different cases give similar values in Equation 27, i.e., populations are not in the true asymptotic regimes corresponding to one case or another. However, since they all roughly agree, the predicted value for 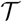 is still accurate.

